# Fast and accurate statistical inference of phylogenetic networks using large-scale genomic sequence data

**DOI:** 10.1101/132795

**Authors:** Hussein A. Hejase, Natalie VandePol, Gregory M. Bonito, Kevin J. Liu

## Abstract

An emerging discovery in phylogenomics is that interspecific gene flow has played a major role in the evolution of many different organisms. To what extent is the Tree of Life not truly a tree reflecting strict “vertical” divergence, but rather a more general graph structure known as a phylogenetic network which also captures “horizontal”gene flow? The answer to this fundamental question not only depends upon densely sampled and divergent genomic sequence data, but also computational methods which are capable of accurately and efficiently inferring phylogenetic networks from large-scale genomic sequence datasets. Recent methodological advances have attempted to address this gap. However, in the 2016 performance study of Hejase and Liu, state-of-the-art methods fell well short of the scalability requirements of existing phylogenomic studies.

The methodological gap remains: how can phylogenetic networks be accurately and efficiently inferred using genomic sequence data involving many dozens or hundreds of taxa? In this study, we address this gap by proposing a new phylogenetic divide-and-conquer method which we call FastNet. We conduct a performance study involving a range of evolutionary scenarios, and we demonstrate that FastNet outperforms state-of-the-art methods in terms of computational efficiency and topological accuracy.

## 1 Introduction

Recent advances in biomolecular sequencing [26] and phylogenomic modeling and inference [8, 30] have revealed that interspecific gene flow has played a major role in the evolution of many different organisms across the Tree of Life [25, 19, 1], including humans and ancient hominins [11, 35], butterflies [40], mice [24], and fungi [10]. These findings point to new directions for phylogenetics and phylogenomics: to what extent is the Tree of Life not truly a tree reflecting strict vertical divergence, but rather a more general graph structure known as a phylogenetic network where reticulation edges and nodes capture gene flow? And what is the evolutionary role of gene flow? In addition to densely sampled and divergent genomic sequence data, one additional ingredient is needed to make progress on these questions: computational methods which are capable of accurately and efficiently inferring phylogenetic networks on large-scale genomic sequence datasets.

Recent methodological advances have attempted to address this gap. Solís-Lemus and Ané proposed SNaQ [38], a new statistical method which seeks to address the computational efficiency of species network inference using a pseudolikelihood approximation. The method of Yu and Nakhleh [41] (referred to here as MPL, which stands for maximum pseudo-likelihood) substitutes pseudolikelihoods in place of the full model likelihoods used by the methods of Yu et al. [44] (referred to here as MLE, which stands for maximum likelihood estimation, and MLE-length, which differ based upon whether or not gene tree branch lengths contribute to model likelihood). Two of us recently conducted a performance study which demonstrated the scalability limits of SNaQ, MPL, MLE, MLE-length, and other state-of-the-art phylogenetic methods in the context of phylogenetic network inference [13]. The scalability of the state of the art falls well short of that required by current phylogenetic studies, where many dozens or hundreds of divergent genomic sequences are common [30]. The most accurate phylogenetic network inference methods performed statistical inference under phylogenomic models [44, 43, 38] that extended the multi-species coalescent model [20, 12]. MPL and SNaQ were among the fastest of these methods while MLE and MLE-length were the most accurate. None of the statistical phylogenomic inference methods completed analyses of datasets with 30 taxa or more after many weeks of CPU runtime – not even the pseudo-likelihood-based methods which were devised to address the scalability limitations of other statistical approaches. The remaining methods fell into two categories: split-based methods [4, 6] and the parsimony-based inference method of Yu et al. [42] (which we refer to as MP in this study). Both categories of methods were faster than the statistical phylogenomic inference methods but less accurate.

The methodological gap remains: how can species networks be accurately and efficiently inferred using large-scale genomic sequence datasets? In this study, we address this question and propose a new method for this problem. We investigate this question in the context of two constraints. We focus on dataset size in terms of the number of taxa and the number of reticulations in the species phylogeny. We note that scalability issues arise due to other dataset features as well, including population-scale allele sampling for each taxon in a study and sequence divergence.

## 2 Methods

One path forward is through the use of divide-and-conquer. The general idea behind divide-and-conquer is to split the full problem into smaller and more closely related subproblems, analyze the subproblems using state-of-the-art phylogenetic network inference methods, and then merge solutions on the subproblems into a solution on the full problem. Viewed this way, divide-and-conquer can be seen as a computational framework that “boosts” the scalability of existing methods (and which is distinct from boosting in the context of machine learning). The advantages of analyzing smaller and more closely related subproblems are two-fold. First, smaller subproblems present more reasonable computational requirements compared to the full problem. Second, the evolutionary divergence of taxa in a subproblem is reduced compared to the full set of taxa, which has been shown to improve accuracy for phylogenetic tree inference [15, 9, 22]. We and others have successfully applied divide-and-conquer approaches to enable scalable inference in the context of species tree estimation [22, 23, 29].

Here, we consider the more general problem of inferring species phylogenies that are directed phylogenetic networks. A directed phylogenetic network *N* = (*V*, *E*) consists of a set of nodes *V* and a set of directed edges *E*. The set of nodes *V* consists of a root node *r*(*N*) with in-degree 0 and out-degree 2, leaves 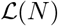 with in-degree 1 and out-degree 0, tree nodes with in-degree 1 and out-degree 2, and reticulation nodes with in-degree 2 and out-degree 1. A directed edge (*u*,*v*) ∈ *E* is a tree edge if and only if *v* is a tree node, and is otherwise a reticulation edge. Following the instantaneous admixture model used by Durand et al. [7], each reticulation node contributes a parameter γ, where one incoming edge has admixture frequency γ and the other has admixture frequency 1−γ. The edges in a network *N* can be labeled by a set of branch lengths ℓ A directed phylogenetic tree is a special case of a directed phylogenetic network which contains no reticulation nodes (and edges). An unrooted tree can be obtained from a directed tree by ignoring edge directionality.

The phylogenetic network inference problem consists of the following. One input is a partitioned multiple sequence alignment ***A*** containing data partitions *a*_*i*_ for 1 ≤ *i* ≤ *k*, where each partition corresponds to the sequence data for one of *k* genomic loci. Each of the *n* rows in the alignment ***A*** is a sample representing taxon *x* ∈ *X*, and each taxon is represented by one or more samples. Similar to other approaches [44, 38], we also require an input parameter *C*_*r*_ which specifies a hypothesized number of reticulations. We note that increasing *C*_*r*_ for a given input alignment ***A*** results in a solution with either better or equal likelihood under the evolutionary models used in our study and others [44, 38]. As is common practice for this and many other statistical inference/learning problems, inference can be coupled with standard model selection techniques (e.g., information criteria [2, 3, 16, 37], cross-validation, etc.) to balance model fit to the observed data against model complexity, thereby determining a suitable choice for parameter *C*_*r*_ in an automated manner. The output consists of a directed phylogenetic network *N* where each leaf in 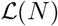 corresponds to a taxon *x* ∈ *X*.

### 2.1 The FastNet algorithm

We now describe our new divide-and-conquer algorithm, which we refer to as FastNet. A flowchart of the algorithm is shown in Figure 1. (Detailed pseudocode can be found in the Appendix’s Supplementary Methods section.)

**Fig. 1.**
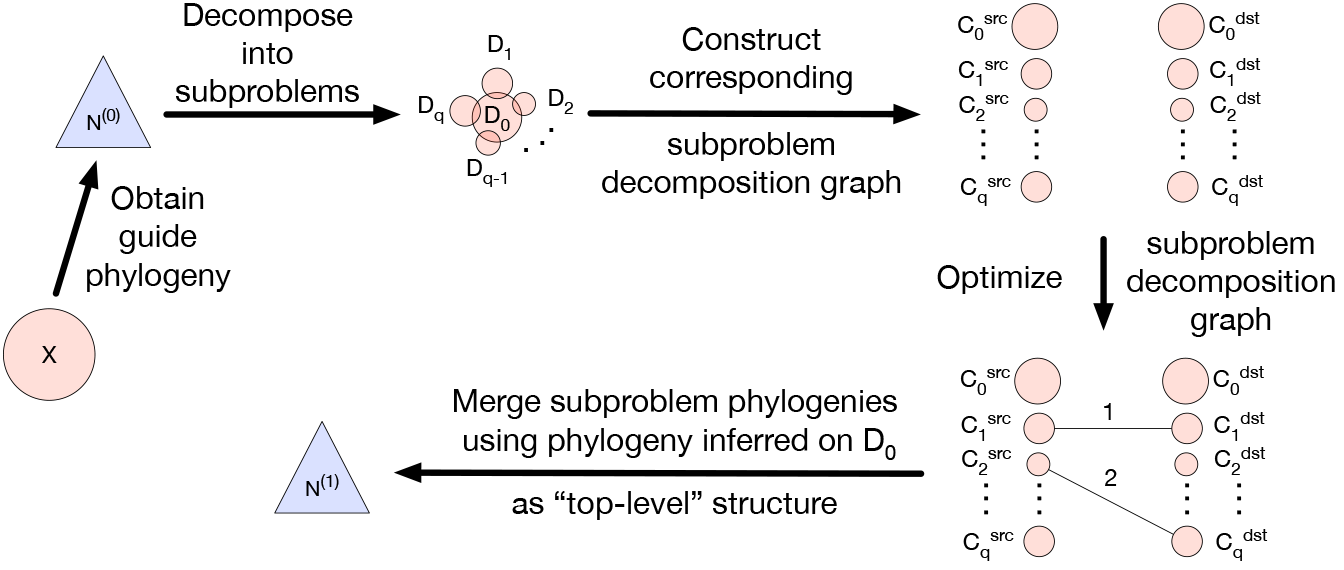
A high-level illustration of the FastNet algorithm. First, a guide phylogeny *N*^(0)^is inferred on the full set of taxa *X*. Next, the guide phylogeny *N*^(0)^is used to decompose *X* into subproblems {*D*_0_, *D*_1_, *D*_2_,…, *D*_*q*−1_, *D*_*q*_} = *D*. Then, the subproblem decomposition *D* is used to construct a bipartite graph *G*_*D*_ = (*V*_*D*_, *E*_*D*_), which is referred to as the subproblem decomposition graph. The set of vertices *V*_*D*_ consist of two partitions: source vertices 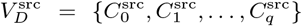 where each subproblem *D*_*i*_ has a corresponding source vertex 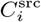, and destination vertices 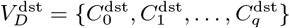 similarly. The subproblem decomposition graph *G*_*D*_ is optimized to infer subproblem phylogenies and reticulations, where the latter are inferred based on the placement of weighted edges *e* ∈ *E*_*D*_. Finally, the subproblem phylogenies are merged using the phylogeny inferred on *D*_0_ as the “top-level” structure.

#### Step zero: obtaining local gene trees

FastNet is a summary-based method for inferring phylogenetic networks. Each subsequent step of the FastNet algorithm therefore utilizes a set of gene trees *G* as input, where a gene tree *g*_*i*_ ∈ *G* represents the evolutionary history of each data partition *a*_*i*_. The experiments in our study utilized either true or inferred gene trees as input to summary-based inference methods, including FastNet (see below for details). We used FastTree [33] to perform maximum likelihood estimation of local gene trees. Our study made use of an outgroup, and the unrooted gene trees inferred by FastTree were rooted on the leaf edge corresponding to the outgroup.

#### Step one: obtaining a guide phylogeny

The subsequent subproblem decomposition step requires a rooted guide phylogeny *N*^(0)^. The phylogenetic relationships need not be completely accurate. Rather, the guide tree needs to be sufficiently accurate to inform subsequent divide-and-conquer steps. Another essential requirement is that the method used for inferring the guide phylogeny must have reasonable computational requirements.

Based on these criteria, we utilized two different methods to obtain guide phylogenies. We used the parsimony-based algorithm proposed by Yu et al. [42] to infer a rooted species network. The algorithm is implemented in the PhyloNet software package [39]. We refer to this method as MP. In a previous simulation study [13], we found that MP offers a significant runtime advantage relative to other state-of-the-art species network inference methods, but had relatively lower topological accuracy. We also used ASTRAL [27, 28], a state-of-the-art phylogenomic inference method that infers species trees, to infer a guide phylogeny that was a tree rather than a network. A primary reason for the use of species tree inference methods is their computational efficiency relative to state-of-the-art phylogenetic network inference methods. While ASTRAL accurately infers species trees for evolutionary scenarios lacking gene flow, the assumption of tree-like evolution is generally invalid for the computational problem that we consider. As we show in our performance study, our divide-and-conquer approach can still be applied despite this limitation, suggesting that FastNet is robust to guide phylogeny error. Another consideration is that ASTRAL effectively infers an unrooted and undirected species tree. We rooted the species tree using outgroup rooting.

#### Step two: subproblem decomposition

The rooted and directed species network *N*^(0)^ is then used to produce a subproblem decomposition D. The decomposition *D* consists of a “bottom-level” component and a “top-level” component, which refers to the subproblem decomposition technique. The bottom-level component is comprised of disjoint subsets *D*_*i*_ for 1 ≤ *i* ≤ *q* which partition the set of taxa *X* such that 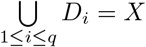. We refer to each subset *D*_*i*_ as a bottom-level subproblem. The top-level component consists of a top-level subproblem *D*_0_ which overlaps each bottom-level subproblem *D*_*i*_ where 1 ≤ *i* ≤ *q*.

The bottom-level component of the subproblem decomposition is obtained using the following steps. First, for each reticulation node in the network *N*^(0)^, we delete the incoming edge with lower admixture frequency. Since the resulting phylogeny *T*^(0)^ contains no reticulation edges and is therefore a tree, removal of any single edge will disconnect the phylogeny into two subtrees; the leaves of the two subtrees will form two subproblems. We extend this observation to obtain decompositions with two or more subproblems. Let *S* be an open set of nodes in the guide phylogeny *T*^(0)^. Each node *s* ∈ *S* induces a corresponding subproblem *D*_*i*_ for 1 ≤ *i* ≤ *q* which consists of the taxa corresponding to the leaves that are reachable from *s* in *T*^(0)^. Of course, not all decompositions are created equal. In this study, we explore the use of two criteria to evaluate decompositions: the maximum subproblem size *c*_*m*_ and a lower bound on the number of subproblems. We addressed the resulting optimization problem using a greedy algorithm. The algorithm is similar to the Center-Tree-*i* decomposition used by Liu et al. [22] in the context of species tree inference. The main difference is that we parameterize our divide-and-conquer based upon a different set of optimization criteria. The input to our decomposition algorithm is the rooted directed tree *T*^(0)^ and the parameter *c*_*m*_, which specifies the maximum subproblem size. Our decomposition procedure also stipulates a minimum number of subproblems of 2. Initially the open set *S* consists of the root node *r*(*T*^(0)^). The open set *S* is iteratively updated as follows: each iteration greedily selects a node *s* ∈ *S* with maximal corresponding subproblem size, the node *s* is removed from the set *S* and replaced by its children. Iteration terminates when both decomposition criteria (the maximum subproblem size criterion and the minimum number of subproblems) are satisfied. If no decomposition satisfies the criteria, then the search is restarted using a maximum subproblem size of *c*_*m*_ − 1. In practice, the parameter *c*_*m*_ is set to an empirically determined value which is based upon the largest datasets that state-of-the-art methods can analyze accurately within a reasonable timeframe [13]. The output of the search algorithm is effectively a search tree 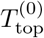 with a root corresponding to *r*(*T*^(0)^), leaves corresponding to *s* ∈ *S*, and the subset of edges in *T*^(0)^ which connect the root *r*(*T*^(0)^) to the nodes *s* ∈ *S* in *T*^(0)^. The decomposition is obtained by deleting the search tree’s corresponding edge structure in *T*^(0)^, resulting in *q* sub-trees which induce bottom-level subproblems as before.

The top-level component augments the subproblem decomposition with a single top-level subproblem *D*_0_ which overlaps each bottom-level subproblem. Phylogenetic structure inferred on *D*_0_ represents ancestral evolutionary relationships among bottom-level subproblems. Furthermore, overlap between the top-level subproblem *D*_0_ and bottom-level subproblems is necessary for the subsequent merge procedure (see “Step four” below). The top-level subproblem *D*_0_ contains representative taxa taken from each bottom-level subproblem *D*_*i*_ for 1 ≤ *i* ≤ *q*: for each bottom-level subproblem *D*_*i*_, we choose the leaf in *T*^(0)^ that is closest to the corresponding open set node *s* ∈ *S* to represent *D*_*i*_, and the corresponding taxon is included in the top-level subproblem *D*_0_.

#### Step three: subproblem decomposition graph optimization

Tree-based divide-and-conquer approaches reduce evolutionary divergence within subproblems by effectively partitioning the inference problem based on phylogenetic relationships. Within each part of the true phylogeny corresponding to a subproblem, the space of possible unrooted sub-tree topologies contributes a smaller set of distinct bipartitions (each corresponding to a possible tree edge) that need to be evaluated during search as compared to the full inference problem. The same insight can be applied to reticulation edges as well, except that a given reticulation is not necessarily restricted to a single subproblem.

We address the issue of “inter-subproblem” reticulations through the use of an abstraction which we refer to as a subproblem decomposition graph. A subproblem decomposition graph *G*_*D*_ = (*V*_*D*_, *E*_*D*_) is a bipartite graph where the vertices *V*_*D*_ can be partitioned into two sets: a set of source vertices 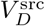 and a set of destination vertices 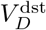. There is a source vertex 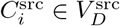 for each distinct subproblem *D*_*i*_ ∈ *D* where 0 ≤ *i* ≤ *q*, and similarly for destination vertices 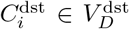. An undirected edge *e*_*ij*_ ∈ *E*_*D*_ connects a source vertex 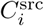 to a destination vertex 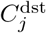 where *i* ≤ *j* and has a weight 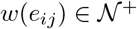. If an edge *e ii* connects nodes 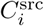 and 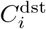 that correspond to the same subproblem *D*_*i*_ ∈ *D*, then the edge weight *w*(*e*_*ii*_) > 0 specifies the number of reticulations in the phylogenetic network to be inferred on subproblem *D*_*i*_; otherwise, a phylogenetic tree is to be inferred on subproblem *D*_*i*_. If an edge *e*_*ij*_ connects nodes 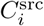 and 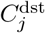 where *i* < *j*, then the edge weight *w*(*e*_*ij*_) > 0 specifies the number of “inter-subproblem” reticulations between the subproblems *D*_*i*_ and *D*_*j*_ (where an intersubproblem reticulation is a reticulation with one incoming edge which is incident from the phylogeny to be inferred on subproblem *D*_*i*_ and the other incoming edge which is incident from the phylogeny to be inferred on *D*_*j*_); otherwise, no reticulations are to be inferred between the two subproblems. A subproblem decomposition graph is constrained to have a total number of reticulations such that 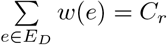.

Given a subproblem decomposition *D*, FastNet’s search routines make use of the correspondence between a subproblem decomposition graph *G*_*D*_ and a multiset with cardinality *C*_*r*_ that is chosen from 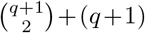 elements. Enumeration over corresponding multisets is feasible when the number of subproblems and *C*_*r*_ are sufficiently small; otherwise, perturbations of a corresponding multiset can be used as part of a local search heuristic. See Algorithm 1 in the Appendix’s Supplementary Methods section for detailed pseudocode.

A subproblem decomposition graph *G*_*D*_ facilitates phylogenetic inference given a subproblem decomposition *D*. The resulting inference is evaluated with respect to a pseudo-likelihood-based criterion. Pseudocode for the pseudo-likelihood calculation is shown in Algorithm 2 in Appendix’s Supplementary Methods section.

The first step is to analyze each individual subproblem *D*_*i*_ ∈ *D* where 0 ≤ *i* ≤ *q*. If an edge *e*_*ii*_ exists, then a phylogenetic network with *w*(*e*_*ii*_) reticulations is inferred on the corresponding subproblem *D*_*i*_; otherwise, a phylogenetic tree is inferred. We utilized MLE, a summary-based MLE method, to perform phylogenetic inference on subproblems, which we refer to as a base method. We note that alternative optimization-based approaches (e.g., other likelihood-based approaches such as MLE-length or pseudo-likelihood-based approaches such as MPL) can be substituted in a straightforward manner.

Next, reticulations are inferred “between” pairs of subproblems as follows. Let *N*_*i*_ and *N*_*j*_ where *i* ≠ *j* be the networks inferred on subproblems *D*_*i*_ and *D*_*j*_, respectively, using the above procedure. Construct the cherry (*N*_*i*_: *b*_*i*_, *N*_*j*_ : *b*_*j*_)ANC; which consists of a new root node ANC with children *r*(*N_i_*) and *r*(*N*_*j*_), where *N*_*i*_ and *N*_*j*_ are respectively retained as sub-phylogenies. Then, infer branch lengths *b*_*i*_ and *b*_*j*_ and add *w*(*e*_*ij*_) reticulations under the maximum likelihood criterion used by the base method. For pairs of subproblems not involving the top-level subproblem *D*_0_, we used the base method to perform constrained optimization. For pairs of subproblems involving the top-level subproblem *D*_0_, we used a greedy heuristic: initial placements were chosen arbitrarily for each reticulation, the source node for each reticulation edge was exhaustively optimized, and then the destination node for each reticulation edge was exhaustively optimized.

Inferred phylogenies and likelihoods were cached to ensure consistency among individual and pairwise subproblem analyses, which is necessary for the subsequent merge procedure. Caching also aids computational efficiency.

Finally, the subproblem decomposition graph and associated phylogenetic inferences are evaluated using a pseudolikelihood criterion:

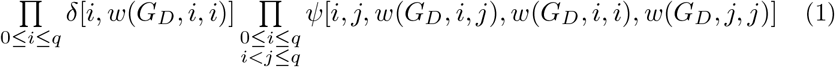

where *w*(*G*_*D*_,*i*,*j*) is the weight of edge *e*_*ij*_ if it exists in *E*(*G*_*D*_) or 0 otherwise, δ[*i*,*w*(*G*_*D*_, *i*, *i*)] is the cached likelihood for an individual subproblem *D*_*i*_, and *ψ*[*i*, *j*,*w*(*G*_*D*_, *i*,*j*),*w*(*G*_*D*_, *i*,*i*),*w*(*G* _*D*_, *j*, *j*)] is the cached likelihood for a pair of subproblems *D*_*i*_ and *D*_*j*_ where *i* < *j*. The pseudo-likelihood calculation effectively assumes that subproblems are independent, although they are correlated through connecting edges in the model phylogeny. The choice of optimization criterion in this context represents a tradeoff between efficiency and accuracy, and several other state-of-the-art phylogenetic inference methods also use pseudolikelihoods to analyze subsets of taxa (e.g., MPL and SNaQ). Other choices are possible. For example, an alternative would be to merge subproblem inferences into a single network hypothesis and calculate its likelihood under the MSNC model.

We optimize subproblem decomposition graphs under the pseudo-likelihood criterion. Exhaustive enumeration of subproblem decomposition graphs is possible for the datasets in our study. Pseudocode to obtain a global optimum is shown in Algorithm 3 in the Appendix’s Supplementary Methods section. For larger datasets with more reticulations, heuristic search techniques can be used to obtain local optima as a more efficient alternative.

#### Step four: merge subproblem phylogenies into a phylogeny on the full set of taxa

Given an optimal subproblem decomposition graph *G*′_*D*_ returned by the previous step, the final step of the FastNet algorithm merges the “top-level” phylogenetic structure inferred on *D*_0_ and “bottom-level” subproblem phylogenies *D*_*i*_ for 1 ≤ *i* ≤ *q* (Algorithm 4 in the Appendix’s Supplementary Methods section). First, the phylogeny inferred on the top-level subproblem *D*_0_ serves as the top-level of the output phylogeny *N′*. Next, the *i*th taxon in *N′* is replaced with the phylogeny inferred on bottom-level subproblem *D*_*i*_, which was cached during the evaluation of *G′*_*D*_. Finally, each “inter-subproblem” reticulation that was inferred for a pair of subproblems *D*_*i*_ and *D*_*j*_ where *i* < *j* is added to the output phylogeny *N′*, which is compatible by construction of the decomposition *D* and the optimal subproblem decomposition graph *G′*_*D*_. The result of the merge procedure is an output phylogeny *N′* on the full set of taxa *X*.

### 2.2 Performance study

We conducted a simulation study to evaluate the performance of FastNet and existing state-of-the-art methods for phylogenetic network inference. The performance study utilized the following procedures. Detailed commands and software options are given in the Supplementary Material (see Additional File 1).

We also conducted an empirical study to evaluate FastNet’s performance. Details about the empirical study are provided in the Appendix.

#### Simulation of model networks

For each model condition, random model networks were generated using the following procedure. First, r8s version 1.7[36] was used to simulate random birth-death trees with *n* taxa where *n* ∈ {15, 20, 25, 30}, which served as in-group taxa during subsequent analysis. The height of each tree was scaled to 5.0 coalescent units. Next, a time-consistent level-*r* rooted phylogenetic network [17] was obtained by adding *r* reticulations to each tree, where *r* ∈ [1,4]. The procedure for adding a reticulation consists of the following steps: based on a consistent timing of events in the tree, (1) choose a time *t*_*M*_ uniformly at random between 0 and the tree height, (2) randomly select two tree edges for which corresponding ancestral populations existed during time interval [*t*_*A*_,*t*_*B*_] such that *t*_*M*_ ∈ [*t*_*A*_,*t*_*B*_], and (3) add a reticulation to connect the pair of tree edges. Finally, an outgroup was added to the resulting network at time 15. 0.

Reticulations in our study have the same interpretation as in the study of Leaché et al. [21]. Gene flow is modeled using an isolation-with-migration model, where each reticulation is modeled as a unidirectional migration event with rate 5.0 during the time interval [*t*_*A*_,*t*_*B*_]. We focus on paraphyletic gene flow as described by Leaché et al.; their study also investigated two other classes of gene flow – isolation-with-migration and ancestral gene flow – both of which involve gene flow between two sister species after divergence. Our simulation study omits these two classes since several existing methods (i.e., MLE and MPL) have issues with identifiability in this context. We note that FastNet makes no assumptions about the type of gene flow to be inferred, and identifiability depends on the model used for inference by FastNet’s base method.

As in the study of Leaché et al., we further classify simulation conditions based on whether gene flow is “non-deep” or “deep” based on topological constraints. Non-deep reticulations involve leaf edges only, and all other reticulations are considered to be deep. Similarly, model conditions with non-deep gene flow have model networks with non-deep reticulations only; all other model conditions include deep reticulations and are referred to as deep.

#### Simulation of local genealogies and DNA sequences

We used ms [14] to simulate local gene trees for independent and identically distributed (i.i.d.) loci under an extended multi-species coalescent model, where reticulations correspond to migration events as described above. Each coalescent simulation sampled one allele per taxon. The primary experiments in our study simulated 1000 gene trees for each random model network. Our study also investigated data requirements of different methods by including additional datasets where either 200 or 100 gene trees were simulated for each random model network.

Sequence evolution was simulated using seq-gen [34], which takes the local genealogies generated by ms as input and simulates sequence evolution along each genealogy under a finite-sites substitution model. Our simulations utilized the Jukes-Cantor substitution model [18]. We simulated 1000 bp per locus, and the resulting multi-locus sequence alignment had a total length of 1000 kb.

#### Replicate datasets

A model condition in our study consisted of fixed values for each of the above model parameters. For each model condition, the simulation procedure was repeated twenty times to generate twenty replicate datasets.

#### Species network inference methods

Our simulation study compared the performance of FastNet against existing methods which were among the fastest and most accurate in our previous performance study of state-of-the-art species network inference methods [13]. Like FastNet, these methods perform summary-based inference – i.e., the input consists of gene trees inferred from sequence alignments for multiple loci, rather than the sequence alignments themselves. The methods are broadly characterized by their statistical optimization criteria: either maximum likelihood or maximum pseudo-likelihood under the multi-species network coalescent (MSNC) model [43]. The maximum likelihood estimation methods consisted of two methods proposed by Yu et al. [44] which are implemented in PhyloNet [39]. One method utilizes gene trees with branch lengths as input observations, whereas the other method considers gene tree topologies only; we refer to the methods as MLE-length and MLE, respectively. Our study also included the pseudo-likelihood-based method of [42], which we refer to as MPL. For each analysis in our study, all species network inference methods – MLE, MLE-length, MPL, and FastNet – were provided with identical inputs.

Our study included two categories of experiments. The “boosting” experiments in our simulation study compared the performance of FastNet against its base method; we refer to all other experiments in our study as “non-boosting”. To make boosting comparisons explicit, each boosting experiment will refer to “FastNet(BaseMethod)” which is FastNet run with a specific base method “BaseMethod” – either MLE-length, MLE, or MPL. The input for each boosting experiment consisted of either true or inferred gene trees for all loci. The inferred gene trees were obtained using FastTree [33] with default settings to perform maximum likelihood estimation under the Jukes-Cantor substitution model [18]. The inferred gene trees were rooted using the outgroup. The non-boosting experiments focused on the performance of FastNet using MLE as a base method and inferred gene trees as input, where gene trees were inferred using the same procedure as in the boosting experiments.

#### Performance measures

The species network inference methods in our study were evaluated using two different criteria. The first criterion was topological accuracy. For each method, we compared the inferred species phylogeny to the model phylogeny using the tripartition distance [31], which counts the proportion of tripartitions that are not shared between the inferred and model network. The second criterion was computational runtime. All computational analyses were run on computing facilities in Michigan State University’s High Performance Computing Center. We used compute nodes in the intel16 cluster, each of which had a 2.5 GHz Intel Xeon E5-2670v2 processor with 64 GiB of main memory. All replicates completed with memory usage less than 16 GiB.

## 3 Results

FastNet’s use of phylogenetic divide-and-conquer is compatible with a range of different methods for inferring rooted species networks on subproblems, which we refer to as “base” methods. From a computational perspective, FastNet can be seen as a general-purpose framework for boosting the performance of base methods. We began by assessing the relative performance boost provided by FastNet when used with two different state-of-the-art network inference methods. We evaluated two different aspects of performance: topological error as measured by the tripartition distance [31] between an inferred species network and the model network, and computational runtime. The initial set of boosting experiments focused on species network inference in isolation of upstream inference accuracy by providing true gene trees as input to all of the summary-based inference methods.

In the performance study of Hejase and Liu [13], the probabilistic network inference methods were found to be the most accurate among state-of-the-art methods, and MPL was among the fastest methods in this class. MPL utilized a pseudo-likelihood-based approximation for increased computational efficiency compared with full likelihood methods [41]. However, the tradeoff netted efficiency that was well short of current phylogenomic dataset sizes [13].

Table 1 shows the performance of FastNet(MPL) relative to MPL on model conditions with increasing numbers of taxa and non-deep reticulations. On model conditions with dataset sizes ranging from 15 to 30 taxa and from 1 to 4 reticulations, FastNet(MPL)’s improvement in topological error relative to its base method was statistically significant (one-sided pairwise t-test with Benjamini-Hochberg correction for multiple tests [5]; *α* = 0.05 and *n* = 20) and substantial in magnitude – an absolute improvement that amounted to as much as 41%. Furthermore, the improvement in topological error grew as datasets became larger and involved more reticulations: the largest improvements were seen on the 30-taxon 4-reticulation model condition. Runtime improvements were also statistically significant and represented speedups which amounted to as much as a day and a half of runtime.

**Table 1.**
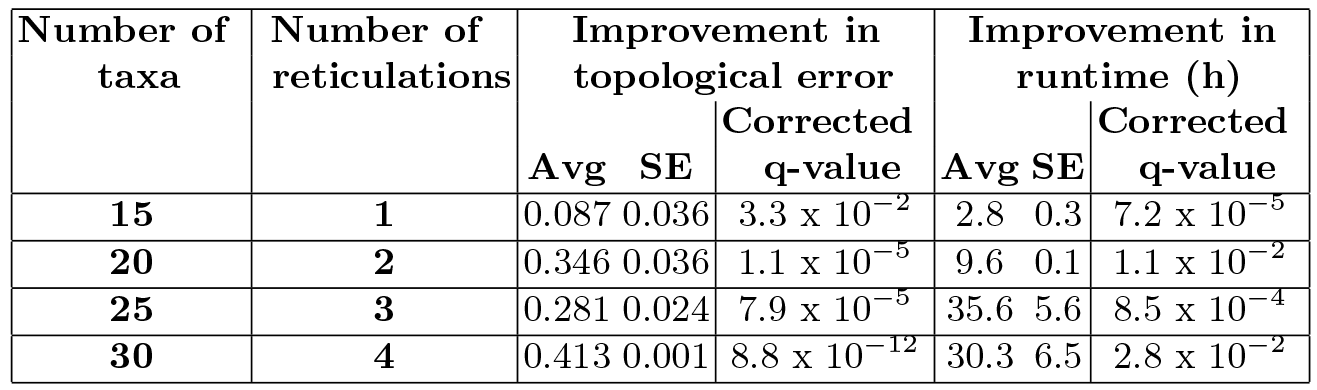
FastNet(MPL) “boosts” MPL’s runtime and topological accuracy, where a greater performance boost occurs as dataset sizes increase. The relative performance of FastNet(MPL) and MPL is compared on model conditions with 15-30 taxa and 1-4 non-deep reticulations. The performance measures consisted of topological error as measured by the tripartition distance between an inferred species network and the model network and computational runtime in hours. Average (“Avg”) and standard error (“SE”) of FastNet(MPL)’s performance improvement over MPL is reported (*n* = 20). All methods were provided with true gene trees as input. The statistical significance of FastNet(MPL)’s performance improvement over MPL was assessed using a one-sided t-test. Corrected q-values are reported where multiple test correction was performed using the Benjamini-Hochberg method [5].

Next, we evaluated FastNet’s performance when boosting MLE-length, the most accurate state-of-the-art method from the performance study of Hejase and Liu [13]. On model conditions with non-deep reticulations, FastNet(MLE-length) had a similar boosting effect as compared to FastNet(MPL) (Table 2). On the 15-taxon single-reticulation model condition, FastNet’s average improvement in topological error was greater when MLE-length was used as a base method rather than MPL. An even greater improvement in computational runtime was seen: FastNet(MLE-length)’s runtime improvement over MLE-length was over an order of magnitude greater than FastNet(MPL)’s improvement over MPL. As the number of taxa increased from 15 to 20 (but the number of reticulations was fixed to one), FastNet(MLE-length)’s advantage in topological error and runtime relative to its base method nearly doubled. In all cases, FastNet(MLE-length)’s performance improvements were statistically significant (Benjamini-Hochberg-corrected one-sided pairwise t-test; *α* = 0.05 and *n* = 20). Although FastNet(MLE-length) successfully completed analysis of larger datasets (i.e., model conditions with more than 20 taxa and/or more than one reticulation), we were unable to quantify FastNet(MLE-length)’s performance relative to its base method due to MLE-length’s scalability limitations.

**Table 2.**
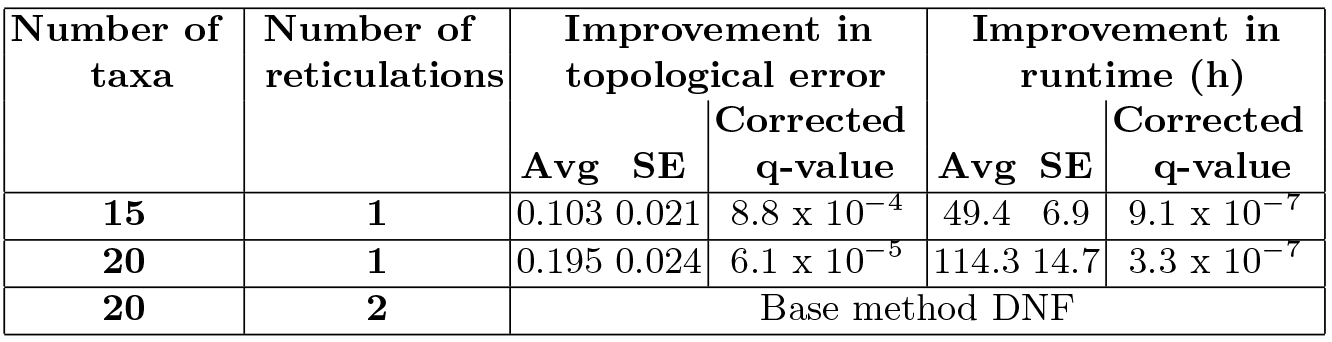
FastNet(MLE-length) “boosts” MLE-length’s runtime and topological accuracy, where a greater performance boost occurs as dataset sizes increase. The relative performance of FastNet(MLE-length) and MLE-length is compared on model conditions with 15-20 taxa and 1-2 non-deep reticulations. Note that, for the model condition with 20 taxa and 2 reticulations, MLE-length did not finish analysis of any replicates after a week of runtime. Otherwise, table layout and description are identical to Table 1.

We further evaluated FastNet’s performance in the context of additional experimental and methodological considerations. On model conditions with deep gene flow (Table 3), FastNet returned significant improvements in topological accuracy and runtime relative to its base method – either MPL or MLE-length – with one exception: on the 15-taxon single-reticulation model condition, Fast Net(MPL) returned a small and statistically insignificant improvement in topological error over MPL. Otherwise, FastNet’s performance boost was robust to the choice of base method. As dataset sizes increased, the average performance boost increased when MPL was the base method; a similar finding applied to runtime improvements when MLE-length was the base method, whereas topological error improvements were largely unchanged. We note that FastNet’s performance boost was somewhat smaller on model conditions involving deep gene flow as opposed to non-deep gene flow. When maximum-likelihood-estimated gene trees were used as input to summary-based inference in lieu of true gene trees (Table 4), FastNet boosted the topological accuracy and runtime of its base method in all cases and the improvements were statistically significant. As dataset sizes increased, FastNet’s improvement in topological accuracy and runtime grew when MPL was its base method; runtime improvements grew and topological error improvements were largely unchanged when MLE-length was the base method. Finally, we conducted an additional experiment to evaluate FastNet’s statistical efficiency when given a finite number of observations in terms of the number of loci (Table 5). As the number of loci ranged from genome-scale (i.e., on the order of 1000 loci) to sizes that were smaller by up to an order of magnitude, FastNet’s average topological error increased by less than 0.02.

**Table 3.**
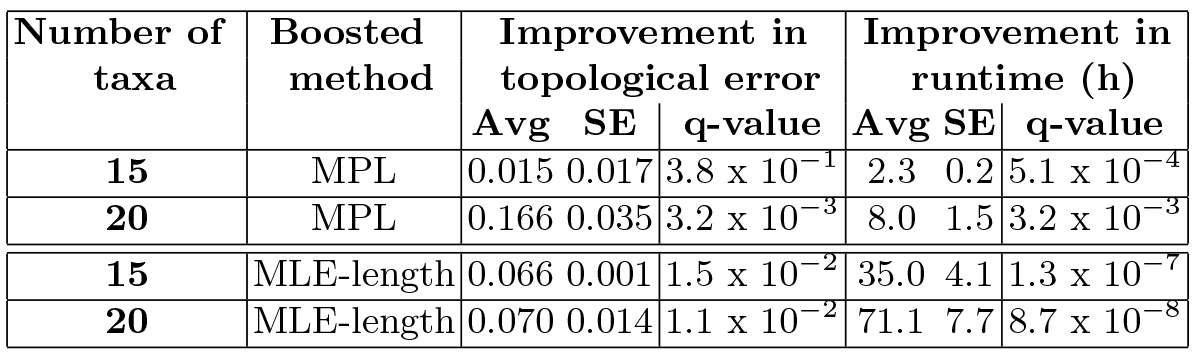
Boosting experiments on model conditions with deep gene flow. The performance improvement of FastNet over its base method (either MPL or MLE-length) is reported for two different performance measures: topological error as measured by tripartition distance and computational runtime in hours. The simulation conditions involved either 15 or 20 taxa and a single deep reticulation. Otherwise, table layout and description are identical to Table 1.

| Number of taxa | Boosted method | Improvement in topological error |  |  | Improvement in runtime (h) |  |  |
| --- | --- | --- | --- | --- | --- | --- | --- |
|  |  | Avg | SE | q-value | Avg | SE | q-value |
| 15 | MPL | 0.015 | 0.017 | $3.8 \times 10^{-1}$ | 2.3 | 0.2 | $5.1 \times 10^{-4}$ |
| 20 | MPL | 0.166 | 0.035 | $3.2 \times 10^{-3}$ | 8.0 | 1.5 | $3.2 \times 10^{-3}$ |
| 15 | MLE-length | 0.066 | 0.001 | $1.5 \times 10^{-2}$ | 35.0 | 4.1 | $1.3 \times 10^{-7}$ |
| 20 | MLE-length | 0.070 | 0.014 | $1.1 \times 10^{-2}$ | 71.1 | 7.7 | $8.7 \times 10^{-8}$ |

**Table 4.** Boosting experiments using inferred gene trees. The performance improvement of FastNet over its base method (either MPL or MLE-length) is reported for two different performance measures: topological error as measured by tripartition distance and computational runtime in hours. For each replicate dataset, all summary-based methods were provided with the same input: a set of rooted gene trees that was inferred using FastTree and outgroup rooting (see Methods section for more details). The simulation conditions involved either 15 or 20 taxa and 1-2 non-deep reticulations. Otherwise, table layout and description are identical to Table 1.

| Number of taxa | Number of reticulations | Boosted method | Improvement in topological error |  |  | Improvement in runtime (h) |  |  |
| --- | --- | --- | --- | --- | --- | --- | --- | --- |
|  |  |  | Avg | SE | q-value | Avg | SE | q-value |
| 15 | 1 | MPL | 0.071 | 0.021 | $1.2 \times 10^{-2}$ | 3.8 | 0.5 | $7.7 \times 10^{-5}$ |
| 20 | 2 | MPL | 0.134 | 0.017 | $1.4 \times 10^{-2}$ | 15.1 | 1.7 | $6.9 \times 10^{-6}$ |
| 15 | 1 | MLE-length | 0.231 | 0.002 | $1.3 \times 10^{-4}$ | 15.4 | 2.0 | $6.7 \times 10^{-7}$ |
| 20 | 1 | MLE-length | 0.195 | 0.005 | $5.8 \times 10^{-5}$ | 43.2 | 7.3 | $1.7 \times 10^{-5}$ |

**Table 5.** The impact of the number of observed loci on FastNet(MLE)’s topological error. The inputs to FastNet(MLE) consisted of gene trees that were inferred using FastTree and outgroup rooting (see Methods section for more details). The simulations sampled between 100 and 1000 loci for a single 20-taxon 1-reticulation model condition involving non-deep gene flow. Topological error was evaluated based upon the tripartition distance between the model phylogeny and the species phylogeny inferred by FastNet(MLE); average (“Avg”) and standard error (“SE”) are shown (*n* = 20).

| Number of loci | Topological error |  |
| --- | --- | --- |
|  | Avg | SE |
| 100 | 0.094 | 0.028 |
| 200 | 0.078 | 0.024 |
| 1000 | 0.075 | 0.027 |

## 4 Discussion

Relative to the state-of-the-art methods that served as base methods, FastNet consistently returned sizeable and statistically significant improvements in topological error and computational runtime across a range of dataset scales and gene flow scenarios. There was only a single experimental condition where comparable error without statistically significant improvements was seen. This exception occurred when FastNet was used to boost a relatively inaccurate base method (MPL) on the smallest dataset sizes in our study and with deep gene flow; even still, large and statistically significant runtime improvements were seen in this case. In contrast, with a more accurate base method (i.e., MLE-length), large and statistically significant performance improvements were seen throughout our simulation study.

FastNet’s boosting effect on topological error and runtime were robust to several different experimental and design factors. The boosting performance obtained using different base methods – one with lower computational requirements but higher topological error relative to a more computationally intensive alternative – suggests that, while accuracy improvements can be obtained even using less accurate subproblem inference, even greater accuracy improvements can be obtained when reasonably accurate subproblem phylogenies can be inferred. We note that the base methods were run in default mode. More intensive search settings for each base method’s optimization procedures may allow a tradeoff between topological accuracy and computational runtime. We stress that our goal was *not* to make specific recommendations about the nuances of running the base methods. Rather, FastNet’s divide-and-conquer framework can be viewed as orthogonal to the specific algorithmic approaches utilized by a base method. In this sense, improvements to the latter accrue to the former in a straightforward and modular manner. Furthermore, FastNet’s performance effect was robust to gene tree error and varying numbers of observed loci.

The biggest performance gains were observed on the largest, most challenging datasets. The findings in our earlier performance study [13] suggest that, given weeks of computational runtime, even the fastest statistical methods (including MPL) would not complete analysis of datasets with more than 50 taxa or so and several reticulations. In comparison to MPL, FastNet(MPL) was faster by more than an order of magnitude on the largest datasets in our study, and we predict that FastNet(MPL) would readily scale to datasets with many dozens of taxa and multiple reticulations.

## 5 Conclusions

In this study, we introduced FastNet, a new computational method for inferring phylogenetic networks from large-scale genomic sequence datasets. FastNet utilizes a divide-and-conquer algorithm to constrain two different aspects of scale: the number of taxa and evolutionary divergence. We evaluated the performance of FastNet in comparison to state-of-the-art phylogenetic network inference methods. We found that FastNet improves upon existing methods in terms of computational efficiency and topological accuracy. On the largest datasets explored in our study, the use of the FastNet algorithm as a boosting framework enabled runtime speedups that were over an order of magnitude faster than standalone analysis using a state-of-the-art method. Furthermore, FastNet returned comparable or typically improved topological accuracy compared to the state-of-the-art-methods that were used as its base method.

Future enhancements to FastNet’s algorithmic design are anticipated to yield additional performance improvements. First, recursive application of phylogenetic divide-and-conquer will allow “load balancing” of subproblem size and divergence. Second, the use of a better guide phylogeny should yield an improved subproblem decomposition and better subproblem decomposition graph optimization. The FastNet-inferred species phylogeny would be a suitable choice in this regard, as demonstrated by the topological error comparisons in our performance study. This insight naturally suggests an iterative approach: the output phylogeny from one iteration of the FastNet algorithm would be used as the guide phylogeny for a subsequent iteration of the algorithm. Iteration would continue until convergence under a suitable criterion (e.g., FastNet’s pseudo-likelihood).

## Acknowledgments

We gratefully acknowledge the following support: NSF grants no. CCF-1565719 (to KJL), CCF-1714417 (to KJL), and DEB-1737898 (to GMB and KJL), BEACON grants (NSF STC Cooperative Agreement DBI-093954) to GMB and KJL, and computing resources provided by MSU HPCC. We would also like to acknowledge Daniel Neafsey for kindly sending us a processed version of the genomic sequence dataset from [32].

## Bibliography

[1] Richard J Abbott and Loren H Rieseberg. Hybrid speciation. In Encyclopaedia of Life Sciences. John Wiley & Sons, Ltd, Hoboken, NJ, USA, 2012.

[2] H. Akaike. Information theory and an extension of the maximum likelihood principle. In Proceedings of the Second International Symposium on Information Theory, pages 267–281, Budapest, Hungary, 1973. Akademiai Kiado.

[3] H. Akaike. A new look at the statistical model identification. IEEE Transactions on Automatic Control, 19(6):716–723, 1974.

[4] Hans-Jürgen Bandelt and Andreas WM Dress. A canonical decomposition theory for metrics on a finite set. Advances in Mathematics, 92(1):47–105, 1992.

[5] Y. Benjamini and Y. Hochberg. Controlling the false discovery rate: A practical and powerful approach to multiple testing. Journal of the Royal Statistical Society Series B (Methodological), 57(1):289–300, 1995.

[6] David Bryant and Vincent Moulton. Neighbor-Net: an agglomerative method for the construction of phylogenetic networks. Molecular Biology and Evolution, 21(2):255–265, 2004.

[7] Eric Y. Durand, Nick Patterson, David Reich, and Montgomery Slatkin. Testing for ancient admixture between closely related populations. Molecular Biology and Evolution, 28(8):2239–2252, 2011.

[8] Scott V Edwards. Is a new and general theory of molecular systematics emerging? Evolution, 63(1):1–19, 2009.

[9] Joseph Felsenstein. Cases in which parsimony or compatibility methods will be positively misleading. Systematic Biology, 27(4):401–410, 1978.

[10] Emile Gluck-Thaler and Jason C Slot. Dimensions of horizontal gene transfer in eukaryotic microbial pathogens. PLoS Pathogens, 11(10):e1005156, 2015.

[11] Richard E. Green, Johannes Krause, Adrian W. Briggs, Tomislav Maricic, Udo Stenzel, Martin Kircher, Nick Patterson, Heng Li, Weiwei Zhai, Markus Hsi-Yang Fritz, Nancy F. Hansen, Eric Y. Durand, Anna-Sapfo Malaspinas, Jeffrey D. Jensen, Tomas Marques-Bonet, Can Alkan, Kay Prüfer, Matthias Meyer, Hernán A. Burbano, Jeffrey M. Good, Rigo Schultz, Ayinuer Aximu-Petri, Anne Butthof, Barbara Höber, Barbara Höffner, Madlen Siegemund, Antje Weihmann, Chad Nusbaum, Eric S. Lander, Carsten Russ, Nathaniel Novod, Jason Affourtit, Michael Egholm, Christine Verna, Pavao Rudan, Dejana Brajkovic, željko Kucan, Ivan Gušic, Vladimir B. Doronichev, Liubov V. Golovanova, Carles Lalueza-Fox, Marco de la Rasilla, Javier Fortea, Antonio Rosas, Ralf W. Schmitz, Philip L. F. Johnson, Evan E. Eichler, Daniel Falush, Ewan Birney, James C. Mullikin, Montgomery Slatkin, Rasmus Nielsen, Janet Kelso, Michael Lachmann, David Reich, and Svante Pääbo. A draft sequence of the Neandertal genome. Science, 328(5979): 710–722, 2010.

[12] Jotun Hein, Mikkel Schierup, and Carsten Wiuf. Gene Genealogies, Variation and Evolution: a Primer in Coalescent Theory. Oxford University Press, Oxford, 2004.

[13] Hussein A Hejase and Kevin J Liu. A scalability study of phylogenetic network inference methods using empirical datasets and simulations involving a single reticulation. BMC Bioinformatics, 17(1):422, 2016.

[14] Richard R. Hudson. Generating samples under a Wright-Fisher neutral model of genetic variation. Bioinformatics, 18(2):337–338, 2002.

[15] John P Huelsenbeck and David M Hillis. Success of phylogenetic methods in the four-taxon case. Systematic Biology, 42(3):247–264, 1993.

[16] Clifford M. Hurvich and Chih-Ling Tsai. Regression and time series model selection in small samples. Biometrika, 76(2):297–307, 1989.

[17] Daniel H Huson, Regula Rupp, and Celine Scornavacca.Phylogenetic Networks: Concepts, Algorithms and Applications. Cambridge University Press, Cambridge, United Kingdom, 2010.

[18] T.H. Jukes and C.R. Cantor. Evolution of Protein Molecules, pages 21–132. Academic Press, New York, NY, USA, 1969.

[19] Patrick J Keeling and Jeffrey D Palmer. Horizontal gene transfer in eukaryotic evolution. Nature Reviews Genetics, 9(8):605–618, 2008.

[20] John Frank Charles Kingman. The coalescent. Stochastic Processes and their Applications, 13(3):235–248, 1982.

[21] Adam D Leaché, Rebecca B Harris, Bruce Rannala, and Ziheng Yang. The influence of gene flow on species tree estimation: a simulation study. Systematic Biology, page syt049, 2013.

[22] K. Liu, S. Raghavan, S. Nelesen, C. R. Linder, and T. Warnow. Rapid and accurate large-scale coestimation of sequence alignments and phylogenetic trees. Science, 324(5934):1561–1564, 2009.

[23] Kevin Liu, Tandy J. Warnow, Mark T. Holder, Serita M. Nelesen, Jiaye Yu, Alexandros P. Stamatakis, and C. Randal Linder. SATé-II: Very fast and accurate simultaneous estimation of multiple sequence alignments and phylogenetic trees. Systematic Biology, 61(1):90–106, 2012.

[24] Kevin J. Liu, Ethan Steinberg, Alexander Yozzo, Ying Song, Michael H. Kohn, and Luay Nakhleh. Interspecific introgressive origin of genomic diversity in the house mouse. Proceedings of the National Academy of Sciences, 112(1):196–201, 2015.

[25] James O McInerney, James A Cotton, and Davide Pisani. The prokaryotic tree of life: past, present…and future?; Trends in Ecology & Evolution, 23 (5):276–281, 2008.

[26] Michael L Metzker. Sequencing technologies - the next generation. Nature Reviews Genetics, 11(1):31–46, 2010.

[27] Siavash Mirarab and Tandy Warnow. ASTRAL-II: coalescent-based species tree estimation with many hundreds of taxa and thousands of genes. Bioinformatics, 31(12):i44–i52, 2015.

[28] Siavash Mirarab, Rezwana Reaz, Md S Bayzid, Theo Zimmermann, M Shel Swenson, and Tandy Warnow. ASTRAL: genome-scale coalescent-based species tree estimation. Bioinformatics, 30(17):i541–i548, 2014.

[29] Siavash Mirarab, Nam Nguyen, Sheng Guo, Li-San Wang, Junhyong Kim, and Tandy Warnow. PASTA: ultra-large multiple sequence alignment for nucleotide and amino-acid sequences. Journal of Computational Biology, 22 (5):377–386, 2015.

[30] Luay Nakhleh. Computational approaches to species phylogeny inference and gene tree reconciliation. Trends in Ecology & Evolution, 28(12):719 – 728, 2013.

[31] Luay Nakhleh, Jerry Sun, Tandy Warnow, C Randal Linder, Bernard ME Moret, and Anna Tholse. Towards the development of computational tools for evaluating phylogenetic network reconstruction methods. In Pacific Symposium on Biocomputing, volume 8, pages 315–326. World Scientific, 2003.

[32] Daniel E. Neafsey, Robert M. Waterhouse, Mohammad R. Abai, Sergey S. Aganezov, Max A. Alekseyev, James E. Allen, James Amon, Bruno Arcà, Peter Arensburger, Gleb Artemov, Lauren A. Assour, Hamidreza Basseri, Aaron Berlin, Bruce W. Birren, Stephanie A. Blandin, Andrew I. Brockman, Thomas R. Burkot, Austin Burt, Clara S. Chan, Cedric Chauve, Joanna C. Chiu, Mikkel Christensen, Carlo Costantini, Victoria L. M. Davidson, Elena Deligianni, Tania Dottorini, Vicky Dritsou, Stacey B. Gabriel, Wamdaogo M. Guelbeogo, Andrew B. Hall, Mira V. Han, Thaung Hlaing, Daniel S. T. Hughes, Adam M. Jenkins, Xiaofang Jiang, Irwin Jungreis, Evdoxia G. Kakani, Maryam Kamali, Petri Kemppainen, Ryan C. Kennedy, Ioannis K. Kirmitzoglou, Lizette L. Koekemoer, Njoroge Laban, Nicholas Langridge, Mara K. N. Lawniczak, Manolis Lirakis, Neil F. Lobo, Ernesto Lowy, Robert M. MacCallum, Chunhong Mao, Gareth Maslen, Charles Mbogo, Jenny McCarthy, Kristin Michel, Sara N. Mitchell, Wendy Moore, Katherine A. Murphy, Anastasia N. Naumenko, Tony Nolan, Eva M. Novoa, Samantha O’Loughlin, Chioma Oringanje, Mohammad A. Oshaghi, Nazzy Pakpour, Philippos A. Papathanos, Ashley N. Peery, Michael Povelones, Anil Prakash, David P. Price, Ashok Rajaraman, Lisa J. Reimer, David C. Rinker, Antonis Rokas, Tanya L. Russell, N’Fale Sagnon, Maria V. Sharakhova, Terrance Shea, Felipe A. Simão, Frederic Simard, Michel A. Slotman, Pradya Somboon, Vladimir Stegniy, Claudio J. Struchiner, Gregg W. C. Thomas, Marta Tojo, Pantelis Topalis, José M. C. Tubio, Maria F. Unger, John Vontas, Catherine Walton, Craig S. Wilding, Judith H. Willis, Yi-Chieh Wu, Guiyun Yan, Evgeny M. Zdobnov, Xiaofan Zhou, Flaminia Catteruccia, George K. Christophides, Frank H. Collins, Robert S. Cornman, Andrea Crisanti, Martin J. Donnelly, Scott J. Emrich, Michael C. Fontaine, William Gelbart, Matthew W. Hahn, Immo A. Hansen, Paul I. Howell, Fotis C. Kafatos, Manolis Kellis, Daniel Lawson, Christos Louis, Shirley Luckhart, Marc A. T. Muskavitch, José M. Ribeiro, Michael A. Riehle, Igor V. Sharakhov, Zhijian Tu, Laurence J. Zwiebel, and Nora J. Besansky. Highly evolvable malaria vectors: The genomes of 16 Anopheles mosquitoes. Science, 347(6217):1258522, 2015.

[33] M. Price, P. Dehal, and A. Arkin. FastTree 2 - approximately maximum-likelihood trees for large alignments. PLoS ONE, 5(3):e9490, March 2010.

[34] A. Rambaut and N. C. Grassly. Seq-Gen: an application for the Monte Carlo simulation of DNA sequence evolution along phylogenetic trees. Computer Applications in the Biosciences, 13:235–238, 1997.

[35] David Reich, Richard E. Green, Martin Kircher, Johannes Krause, Nick Patterson, Eric Y. Durand, Bence Viola, Adrian W. Briggs, Udo Stenzel, Philip L. F. Johnson, Tomislav Maricic, Jeffrey M. Good, Tomas Marques-Bonet, Can Alkan, Qiaomei Fu, Swapan Mallick, Heng Li, Matthias Meyer, Evan E. Eichler, Mark Stoneking, Michael Richards, Sahra Talamo, Michael V. Shunkov, Anatoli P. Derevianko, Jean-Jacques Hublin, Janet Kelso, Montgomery Slatkin, and Svante Paabo. Genetic history of an archaic hominin group from Denisova Cave in Siberia. Nature, 468(7327):1053–1060, 2010.

[36] Michael J Sanderson. r8s: inferring absolute rates of molecular evolution and divergence times in the absence of a molecular clock. Bioinformatics, 19(2):301–302, 2003.

[37] Gideon Schwarz. Estimating the dimension of a model. The Annals of Statistics, 6(2):461–464, 1978.

[38] Claudia Solís-Lemus and Cécile Ané. Inferring phylogenetic networks with maximum pseudolikelihood under incomplete lineage sorting. PLoS Genetics, 12(3):1–21, 03 2016.

[39] Cuong Than, Derek Ruths, and Luay Nakhleh. PhyloNet: a software package for analyzing and reconstructing reticulate evolutionary relationships. BMC Bioinformatics, 9(1):322, 2008.

[40] The Heliconious Genome Consortium. Butterfly genome reveals promiscuous exchange of mimicry adaptations among species. Nature, 487(7405): 94–98, 2012.

[41] Yun Yu and Luay Nakhleh. A maximum pseudo-likelihood approach for phylogenetic networks. BMC Genomics, 16(Suppl 10):S10, 2015.

[42] Yun Yu, Cuong Than, James H Degnan, and Luay Nakhleh. Coalescent histories on phylogenetic networks and detection of hybridization despite incomplete lineage sorting. Systematic Biology, 60(2):138–149, 2011.

[43] Yun Yu, James H. Degnan, and Luay Nakhleh. The probability of a gene tree topology within a phylogenetic network with applications to hybridization detection. PLoS Genetics, 8(4):e1002660, 04 2012.

[44] Yun Yu, Jianrong Dong, Kevin J Liu, and Luay Nakhleh. Maximum likelihood inference of reticulate evolutionary histories. Proceedings of the National Academy of Sciences, 111(46):16448–16453, 2014.

